# Seasonal influenza vaccine strain selection by quantifying viral fitness

**DOI:** 10.64898/2026.09.15.751745

**Authors:** Luyao Qin, Mengyi Zhang, Xiaoyuan Liu, Xiao Ding, Qianqian Li, Lei Yang, Yifan Qi, Jingze Liu, Hangyu Zhou, Zhiming Li, Wenping Xie, Zichen Li, Yun Ma, Jingqi Yang, Haoyang Wang, Jianwei Wang, Taijiao Jiang, Dayan Wang, Youchun Wang, Aiping Wu

**Affiliations:** State Key Laboratory of Common Mechanism Research for Major Diseases, Suzhou Institute of Systems Medicine, Chinese Academy of Medical Sciences & Peking Union Medical College, Suzhou, Jiangsu, China; Key Laboratory of Pathogen Infection Prevention and Control (Peking Union Medical College), Ministry of Education; State Key Laboratory of Respiratory Health and Multimorbidity, Institute of Medical Biology, Chinese Academy of Medical Science & Peking Union Medical College, Kunming, China; National Institute for Viral Disease Control and Prevention, Chinese Center for Disease Control and Prevention, Beijing, China; College of Life Science and Technology, Huazhong University of Science and Technology, Wuhan, China; Guangzhou National Laboratory, Guangzhou, China; NHC Key Laboratory of Systems Biology of Pathogens, Christophe Mérieux Laboratory, National Institute of Pathogen Biology, Chinese Academy of Medical Sciences & Peking Union Medical College, Beijing, China; National Key Laboratory of Immunity and Inflammation, Suzhou Institute of Systems Medicine, Chinese Academy of Medical Sciences, Suzhou, China; Chinese Center for Disease Control and Prevention & Chinese Academy of Preventive Medicine, Beijing, China

## Abstract

The long-standing challenge of seasonal influenza vaccines providing poor protection is attributed to the rapid and continuous evolution of the virus. In theory, the recommendation of vaccine strain is to predict the dominant strain with best fitness in the upcoming season. Here, we develop FutureFlu, a biologically-grounded framework that quantifies the fitness score of influenza variants in upcoming seasons, by integrating metrics on three levels: molecular genetic divergence, individual immune escape, and population-scale transmission dynamics. The fitness scores of influenza variants present strong positive correlation with their actual observed frequencies in the next seasons. Validation across 24 seasons of three influenza subtypes shows FutureFlu recommends antigenically matched vaccine strains more frequently than annual recommendations, particularly for challenging subtypes: 83.3% versus 45.8% seasons for A/H3N2, and 75.0% versus 33.3% seasons for B/Victoria. Furthermore, FutureFlu significantly outperforms the currently used methods in vaccine strain selection whether or not there is available antigenic data from hemagglutination inhibition (HI) assay, which provides a valuable supplement for WHO vaccine recommendation. To support global public health implementation, an open online platform (futureflu.com.cn) has been established to offer real-time viral fitness predictions and vaccine recommendations.

## Main Text

Seasonal influenza viruses present enormous challenges in disease control due to their rapid and continuous evolution. The observed evolution is conceptually shaped by a complex interaction of genetic variations, host immune response, and epidemiology(*1-3*). Novel variants carrying characteristic mutations in antigenic proteins emerge constantly. Those with enhanced immune escape capacity in host individuals and superior transmission efficiency in the population often exhibit increased fitness, subsequently dominating circulation(*4*). Therefore, the turnover of dominant influenza variants continuously weakens the protective efficacy of existing vaccines, necessitating regular vaccine updates(*5, 6*). In theory, if we can quantify the viral fitness, then we can assess the probability that a variant will become dominant in the upcoming seasons, and select the best suitable vaccine strains to improve the vaccine effectiveness.

Previous studies on viral fitness can be broadly classified into two approaches. The first approach focuses on modeling fitness across different components of phylogenetic trees, from clades and strains to mutations(*7-10*). While these methods provide valuable predictions through statistical correlations, they often lack mechanistic explanations for viral adaptive evolution. The second approach employs explicit evolutionary hypotheses to predict future viral populations. For example, Łuksza et al. defines fitness as functional trade-offs between receptor binding and antibody escape(*11*), which emphasizes that viruses should maintain basic functions while acquiring new traits. However, most laboratory-confirmed gain-of-function mutations fail to circulate in the population(*12*), suggesting that viral fitness emerges from the interplay of multiple biological scales, requiring an comprehensive framework for more accurate quantification.

In real-world influenza virus evolution, diverse evolutionary trajectories are observed before variants achieve dominance, which is widely recognized as the comprehensive outcome of genetic evolution, immune responses, and epidemiological processes(*1-3, 13*). At the molecular level, while most mutations occur as random sporadic events, certain mutations and their combinations can become fixed in the genome(*14*). Subsequently, successful clades diversify from their ancestral strain and differentiate themselves from other coexisting clades. At the individual level, mutations in antigenic epitope drive viruses to escape host immunity established through prior infections or vaccination(*15*). Such immune escape can be evaluated through experimental methods like pseudovirus assays(*16*) and deep mutational scanning (DMS)(*12*), but these approaches are costly and retrospective. Computational models like MLAEP(*17*), E2VD(*18*) and EVEscape(*19, 20*) enable rapid prediction of immune escape(*21*), with EVEscape requiring only sequence and structural data when experimental data are unavailable. At the population level, variant with enhanced human-to-human transmission can establish stable transmission chains and gradually replace previous dominant variant(*22*). This replacement process depends on the variant’s transmission efficiency relative to co-circulating strains, reflecting its competitive advantage in the population(*23*). While molecular divergence, individual escape, and population transmission dynamics have been recognized as key drivers in viral evolution, a comprehensive framework integrating these factors to characterize viral evolutionary fitness remains absent.

Here, we proposed a theoretical framework that integrates molecular genetic divergence, individual immune escape, and population-scale transmission dynamics into a unified, quantitative measure of viral fitness. By linking microscopic sequence changes to macroscopic epidemiological impact, it establishes a new paradigm for quantifying viral fitness landscape, which supports evidence-based vaccine strain selection. This strategy is applied to seasonal influenza virus, termed FutureFlu, which predicts dominant variants in the upcoming influenza seasons and recommends vaccine strains. FutureFlu supports two practical applications for vaccine strain recommendation: (1) prioritizing predicted dominant variants when antigenic assay data are unavailable, and (2) selecting cultured strains that are antigenically closest to predicted dominant variants when such antigenic assay exist. Validation results indicate FutureFlu recommends antigenically matched vaccine strains more frequently than annual recommendations, particularly for the challenging subtypes: 83.3% versus 45.8% seasons for A/H3N2, and 75.0% versus 33.3% seasons for B/Victoria. These results demonstrate robust predictive performance over multiple decades and a meaningful reduction in vaccine-mismatch risk. To support global public health implementation, we have launched an open online platform (https://futureflu.com.cn) providing real-time viral fitness predictions and vaccine strain recommendations. This innovative method enables early warning of potentially dominant variants, providing practical guidance for vaccine strain recommendation to improve vaccine effectiveness.

### FutureFlu quantifies the fitness of influenza virus variant

The evolutionary trajectories of influenza clades before dominance are heterogeneous, suggesting that virus evolution is driven by multiple drivers (fig. S1). Here, we decompose the dynamic evolutionary process of variants from emergence to dominance into three levels (Fig. 1A). First is the genetic divergence at molecular level. Multiple mutations in ancestral viral genome make genetic diversity among descendant strains. While most mutations occur randomly, certain mutations and their combinations can become fixed in the genome, leading to the emergence of novel clades. Second is the immune escape at individual level. Key mutations in surface antigenic proteins enable variants to evade recognition and neutralization of existing antibodies. Third is the transmission at population level, where variants with enhanced transmissibility can rapidly establish dominance through human-to-human transmission. This multiscale framework demonstrates that viral fitness is shaped by molecular evolution, immune escape, and transmission dynamics.

**Fig. 1.**
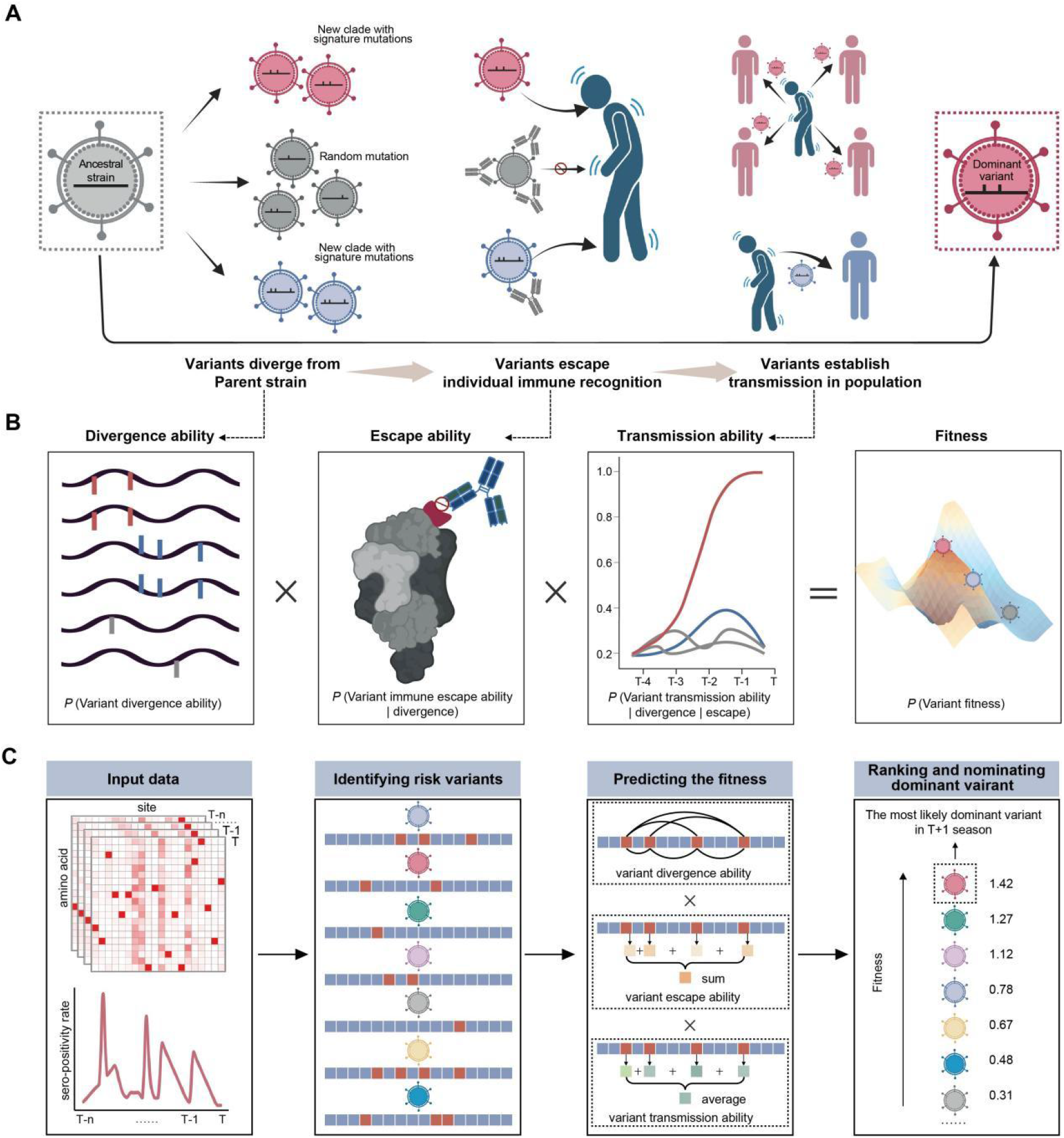
Theoretical framework of viral adaptive evolution and fitness quantification. (**A**) Three levels of selection during virus evolution from emergence to dominance. At the molecular level, genetic diversity arises through multiple mutations in the ancestral genome, where selective mutations become fixed in viral populations, generating distinct independent clades. At the individual level, antigenic drift in surface proteins allows viral variants to escape antibody-mediated immune responses developed from previous exposure. At the population level, variants with enhanced transmissibility can replace existing strains and become dominant in the human population. (**B**) Fitness is defined as the product of three components: divergence ability, escape ability, and transmission ability. (**C**) FutureFlu workflow for predicting the potential dominant variants and their fitness in the upcoming season. The pipeline integrates HA1 amino acid prevalence data and epidemiological sero-positivity rates to identify variants with characteristic mutations (red squares). T denotes the current season, with T-1, T-n representing previous seasons and T+1 indicating the upcoming season. The fitness is quantified by integrating three key components: divergence measured by mutual information among mutations, immune escape assessed by sum of mutation escape scores, and transmission ability derived from the average prevalence of mutations. Higher fitness scores indicate greater likelihood to be dominant in upcoming seasons. Panel (**A**) and (**B**) are created in BioRender.com.

Therefore, we develop a computational framework, FutureFlu, to express the probability that a variant will become dominant in the upcoming seasons, namely fitness score as the product of three probabilities: the likelihoods that a variant will become an independent clade/lineage diverging from the ancestral strain (divergence term), escape from individual immune recognition (escape term), and establish efficient transmission in population (transmission term) (Fig. 1B, fig. S2). In divergence calculation, we employ mutual information to capture non-linear dependencies between co-occurring mutations that characterize independent clades(*24-26*). In escape calculation, we utilize EVEscape, a deep learning model validated across multiple viruses including influenza, to assess antigenic differences between variant and circulating strain using sequence and structure data(*20*). In transmission calculation, we linearly extrapolate the future prevalence of mutations and take their average as the prevalence of variants(*10*). Each component is independently standardized and transformed by temperature-scaled logistic functions to calculate variant fitness while maintaining modularity and interpretability (fig. S3).

It takes HA1 (Hemagglutinin subunit 1) amino acid sequences and epidemiological sero-positivity rate as input. Since viral fitness is enhanced through accumulating advantageous amino acid mutations, leading to more widespread human infections and higher sero-positivity rates, crucial mutations are first identified by analyzing the relationship between cumulative mutation effects and actual sero-positivity rates(*27*). Subsequently, FutureFlu identifies risk variants carrying these crucial mutations and calculates their fitness scores, with higher scores indicating greater potential for dominant circulation in the next season (Fig. 1C).

### FutureFlu predicts dominant variants in next influenza seasons

The World Health Organization (WHO) recommends vaccine strains for the upcoming influenza seasons in February and September for the Northern and Southern Hemispheres (NH and SH), respectively. We conducted retrospective predictions for three subtypes (A/H3N2, A/H1N1(pdm09), and B/Victoria) across 24 influenza seasons from year 2013 to 2024 (including 2013SH-2024SH and 2013NH-2024NH). Due to practical constraints such as sample transportation delays and processing backlogs, the number of available sequences is typically less than that of collected samples(*28*). Therefore, in the retrospective analyses, we only included the GISAID sequences submitted before August 31^st^ or January 31^st^ for SH and NH predictions respectively, to mirror real-world data availability for WHO (Fig. 2A).

**Fig. 2.**
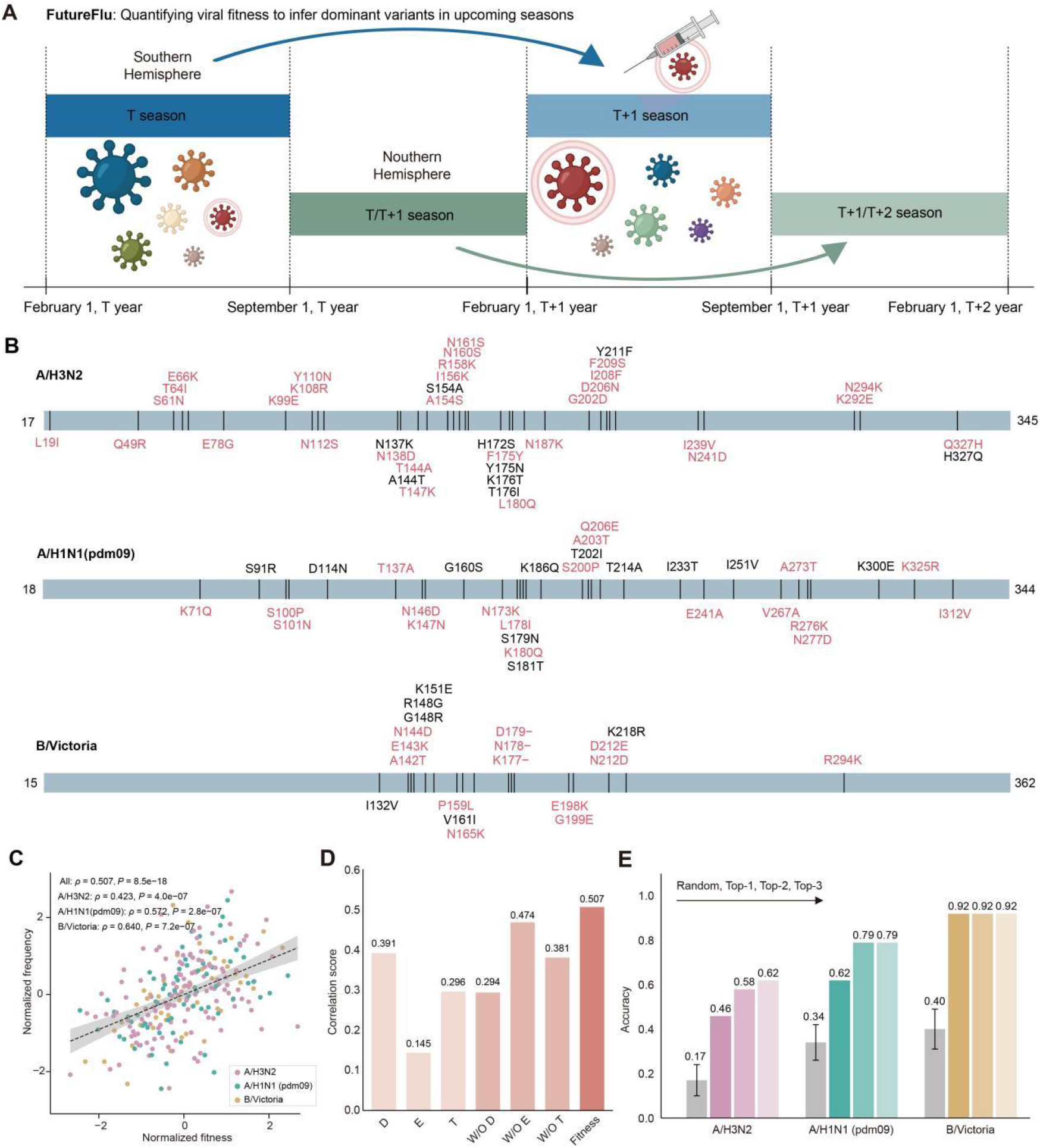
FutureFlu retrospectively predicts variant fitness for the next influenza season. (**A**) The retrospective predictions for dominant variants in upcoming seasons, following WHO’s biannual (February and September) vaccine strain recommendation timeline. This study defines T influenza season in SH as February 1^st^ to August 31^st^ of year T, and T/T+1 influenza season in NH as September 1^st^ of year T to January 31^st^ of year T+1. HA sequences submitted to GISAID up to August 31^st^ of year T are downloaded for prediction of T+1 influenza season in SH. Similarly, HA sequences submitted to GISAID up to January 31^st^ of year T+1 are downloaded for prediction of T+1/T+2 influenza season in NH. The size of the virus indicates its prevalence, with different colors distinguishing various variants. (**B**) Ability of FutureFlu to capture dominant mutations at least one influenza season in advance. Dominant mutations for influenza season T are defined as follows: at a certain position S, when the consensus amino acid of influenza season T (AA_T_) differs from the consensus amino acid of influenza season T-1 (AA_T-1_), and the frequency of AA_T_ in influenza season T significantly increases compared to influenza season T-1 (chi-square test, p<0.05), [AA_T-1_][S][AA_T_] is identified as a dominant mutation for influenza season T. Dominant mutations on the HA1 domain for A/H3N2, A/H1N1(pdm09), and B/Victoria subtypes across 24 seasons are marked as vertical black lines. Gray rectangles represent HA1 domain. Position numbers are sequentially numbered starting from the first amino acid of the HA protein reference sequence for each subtype. Dominant mutations marked in red were captured by FutureFlu at least one influenza season ahead. (**C**) Correlation between variant fitness score and observed frequency. Fitness scores and observed frequencies in each influenza season were quantile-normalized. Pink, green, and yellow dots represent A/H3N2, A/H1N1(pdm09), and B/Victoria, respectively. (**D**) Ablation experiments. Divergence, escape, and transmission are denoted as D, E, and T, respectively. W/O denotes “without”. (**E**) Top-N (N=1, 2, 3) prediction accuracy for dominant variants in 24 seasons of A/H3N2, A/H1N1, and B/Victoria. Gray bars represent mean and standard deviation from 1000 random predictions. Panel **a** is created in BioRender.com.

When predicting at least one season ahead, FutureFlu successfully captured 30/39, 19/30, and 13/19 of actual dominant mutations in A/H3N2, A/H1N1(pdm09), and B/Victoria subtypes, respectively (Fig. 2B). Among the uncaptured dominant mutations, nearly 50% were completely unobserved in earlier surveillance data but emerged and became dominant rapidly (fig. S4a). Additionally, 25% mutations showed high observation frequencies in earlier periods, but underwent multiple amino acid reversions (fig. S4b). These reversion patterns often occurred at positions with multiple functions, reflecting functional trade-offs. For instance, position 214 (H1 numbering from methionine) in A/H1N1(pdm09) HA, located within both the Sb antigenic epitope and receptor binding region, underwent A>T>A substitutions, highlighting the interplay between immune selection and receptor binding(*29*). Similarly, position 154 (H3 numbering from methionine) in A/H3N2 HA, situated in antigenic epitope A, underwent A>S>A substitutions. While these changes affected antigenic recognition, serine at this position also enhanced affinity for the lower respiratory tract and alveolar macrophages in pigs, suggesting this reversion was driven by immune escape and cross-species adaptation at least(*30*).

Variant fitness scores predicted by FutureFlu show significant positive correlation with their actual observed frequencies in next seasons (Spearman correlation, *ρ* = 0.507, *P* < 0.0001) (Fig. 2C). This correlation varies across subtypes, with A/H3N2, A/H1N1(pdm09), and B/Victoria showing correlations of *ρ* = 0.423 (*P* < 0.0001), *ρ* = 0.572 (*P* < 0.0001), and *ρ* = 0.640 (*P* < 0.0001), respectively, reflecting their distinct evolutionary dynamics. Ablation experiments (Fig. 2D, fig. S5) evaluated the contribution of each component and our strategy of integrating multidimensional information. Individual use of divergence (D), escape (E), or transmission (T) terms yielded correlations of ρ = 0.391, 0.145, and 0.296, respectively. Removing any single component (denoted as W/O D, W/O E, or W/O T) resulted in correlation coefficients of ρ = 0.294, 0.474, and 0.381, respectively. For practical application, we evaluated the model’s ability to predict dominant variants. FutureFlu achieved Top-1 prediction accuracies of 46%, 62%, and 92% in A/H3N2, A/H1N1(pdm09), and B/Victoria subtypes, respectively. The prediction accuracy for A/H3N2 and A/H1N1(pdm09) improved when expanding to Top-N predictions (Fig. 2E), indicating that FutureFlu can effectively narrow down the candidate pool of dominant variants even for rapidly evolving viruses.

### FutureFlu selects vaccine strains from the predicted dominant clades

Current vaccine strain selection relies on time-consuming antigenic assays. Based on FutureFlu predictions, we can select vaccine strains from the highest-fitness clade when antigenic assays are unavailable. This approach is effective because antigenic distances between strains within the same clade are significantly smaller than those between strains from different clades (fig. S6). Compared with historical WHO recommendations, LBI(*7*), and beth-1(*10*), FutureFlu achieved higher accuracy in selecting vaccine strains matching dominant clades (Fig. 3A, table S1). For A/H3N2 subtype, FutureFlu vaccine strains matched dominant clades in 11/24 influenza seasons, outperforming WHO recommendations (5/24), LBI (6/24), and beth-1 (5/24). For A/H1N1(pdm09), FutureFlu and beth-1 matched dominant clades in 15/24 seasons, surpassing WHO recommendations (11/24) and LBI (12/24). For B/Victoria, FutureFlu matched dominant clades in 22/24 seasons, outperforming WHO recommendations (18/24), LBI (16/24), and beth-1 (17/24). All methods failed to match dominant clades in some seasons: (1) in 9 seasons, the dominant clades ranked within FutureFlu’s Top-3 predictions (triangles, fig. S7a), (2) in 4 seasons, the dominant clade strain was observed only 1-5 times in available data (squares, fig. S7b), (3) in 9 seasons, the dominant clade strain was not observed in available data (pentagons, fig. S7c), making prediction impossible as the model can only assess the fitness of observed variants.

**Fig. 3.**
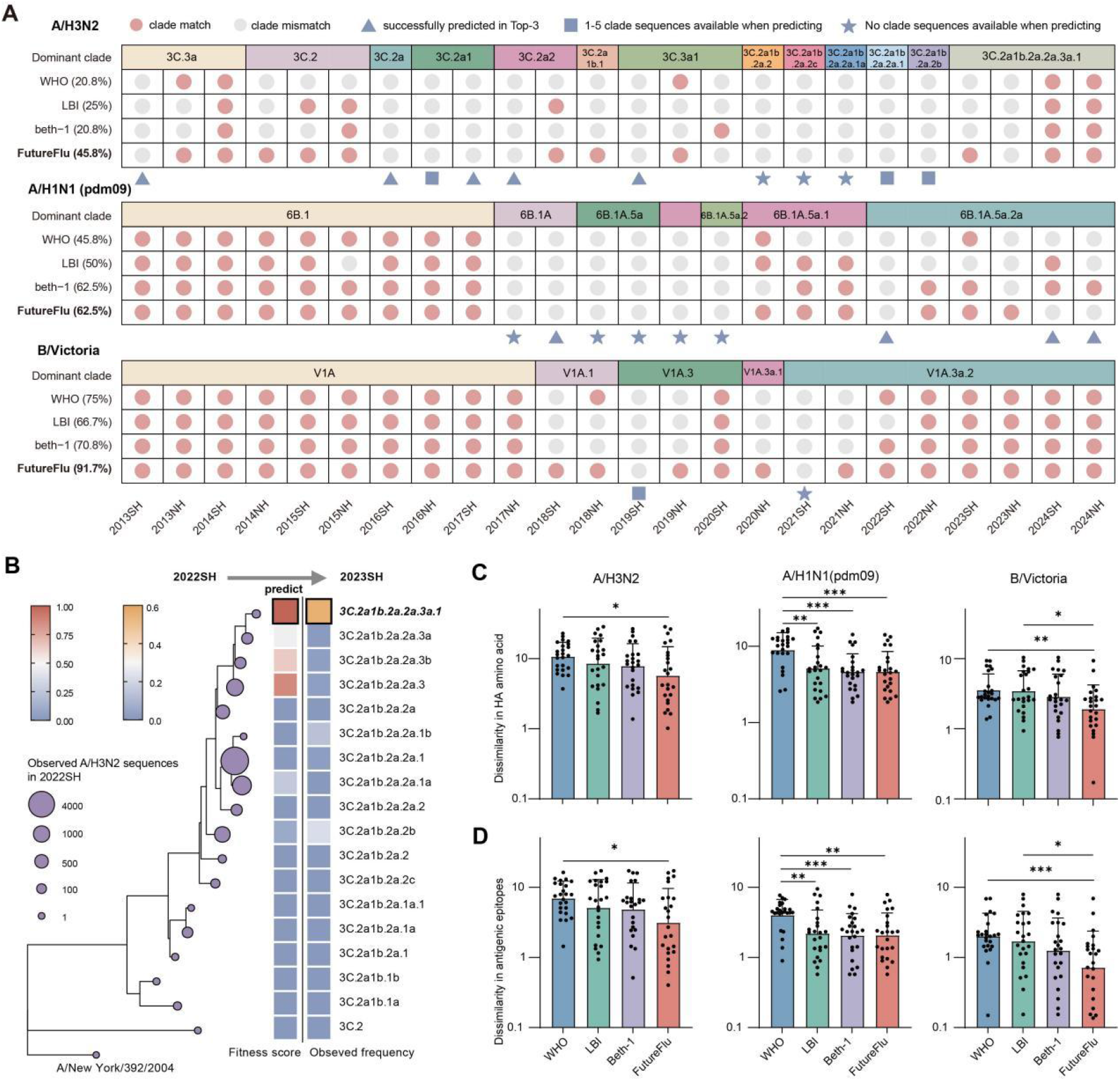
FutureFlu helps to select the vaccine strains matching dominant clades. (**A**) **Matches between vaccine strains and dominant clades in each season**. The dominant clade for each season and vaccine strains recommended by WHO, LBI, beth-1, and FutureFlu are shown in the grid, where red circles indicate successful matches with dominant clades, while gray circles represent mismatches. Blue triangles indicate the dominant clade is among Top-3 predictions of FutureFlu, blue squares indicate that only 1-5 sequences from dominant clades were observed in available data, and blue stars indicate that no sequences from dominant clades were observed in available data. (**B**) FutureFlu captures the dominant clade transition from 2022SH to 2023SH. HA protein sequences submitted to GISAID database up to August 31^st^, 2022 were downloaded for prediction. The representative strain with the highest sampling frequency in each clade was selected for phylogenetic tree construction using FastTree 2.1.11 SSE3. Node size of the phylogenetic tree represents the observed count of clade in 2022SH season. The left heatmap shows the normalized fitness scores of clades, with redder colors indicating higher fitness scores in the 2023SH season. The right heatmap displays the observed frequencies of clades in the 2023SH season, with yellow intensity corresponding to higher frequencies. Black box highlights the clade with highest predicted fitness and the clade with highest observed frequencies. (**C**) Amino acid mismatches at HA protein between vaccine strains and dominant clade strains. Blue, green, purple, and red columns represent vaccine strains in 24 seasons recommended by WHO, LBI, beth-1, and FutureFlu respectively. Each point represents the average number of mismatches between the vaccine strain and dominant clade viruses for a single influenza season. (**D**) Amino acid mismatches at key antigenic epitopes of HA protein between vaccine strains (WHO, LBI, beth-1, FutureFlu) and dominant clade strains.

The advantage of FutureFlu was particularly evident in predicting dominant clade transitions, such as from 2013NH to 2014NH, 2017NH to 2018NH, and 2022SH to 2023SH seasons in A/H3N2 subtype. Specifically, during the 2022SH influenza season, while clade 3C.2a1b.2a.2a.1 was dominant with nearly 20 minor clades co-circulating, FutureFlu correctly predicted that 3C.2a1b.2a.2a.3a.1 has the highest fitness score in 2023SH, which was validated as clade 3C.2a1b.2a.2a.3a.1 indeed became dominant in 2023SH (Fig. 3B). Besides, compared to LBI and beth-1, FutureFlu-recommended strains showed fewer amino acid mismatches in both the entire HA protein (Fig. 3C) and antigenic sites (Fig. 3D). Although influenza circulation patterns exhibit geographic heterogeneity, FutureFlu can assist in predicting influenza evolution trends at the country level. The retrospective analyses for China and United States showed that FutureFlu could recommend vaccine strains matching the dominant clades for more seasons (fig. S8A, B) and exhibited fewer amino acid differences at antigenic sites (fig. S8C, D). These results demonstrate that FutureFlu helps to provide region-specific vaccine recommendations.

### FutureFlu recommends vaccine strains antigenically similar to dominant clades

If antigenic assays, such as Hemagglutination inhibition (HI) assays are available, the cultured strain (from eggs or MDCK cells) with minimal antigenic distance to the predicted dominant clade (with best fitness) is recommended as the vaccine strain (Fig. 4A, table S1). Lower antigenic distances indicate higher vaccine protection efficacy (Fig. 4B). Defining antigenic similarity as distances below 4, FutureFlu recommended antigenically similar cultured strains in more seasons across all subtypes compared to historical WHO recommendations: 20 (83.3%) versus 11 (45.8%) seasons for A/H3N2 (Fig. 4C), 22 (91.7%) versus 21 (87.5%) seasons for A/H1N1 (Fig. 4D), and 18 (75.0%) versus 8 (33.3%) seasons for B/Victoria (fig. S9). Additionally, FutureFlu recommendations have smaller antigenic distances to dominant clades than historical WHO recommendations, with significant improvement for A/H3N2 and B/Victoria (A/H3N2: 2.65 ± 1.70 *vs*. 10.59 ± 11.73, *P* < 0.01; A/H1N1(pdm09): 3.30 ± 8.59 *vs*. 4.48 ± 8.50, *P* > 0.05; B/Victoria: 3.36 ± 4.79 *vs*. 13.50 ± 15.90, *P* < 0.01). During the 10 influenza seasons from year 2013 to 2017, although the WHO recommendation B/Brisbane/60/2008 belonged to V1A, which was the dominant clade, antigenic drift occurred between the early V1A and later circulating V1A strains during long-term evolution (fig. S6B). This resulted in B/Brisbane/60/2008 being unable to protect against the circulating strains. Therefore, the timeliness of vaccine strains should also be considered, avoiding the use of historical strains from too early periods. In comparison with VaxSeer(*31*) recommendations from 2013NH to 2021NH seasons, FutureFlu recommended antigenically similar A/H3N2 vaccine strains for 7/9 seasons, surpassing VaxSeer (5/9), and had the same performance as VaxSeer for A/H1N1(pdm09) (both 9/9 seasons). Moreover, FutureFlu recommendations exhibited smaller antigenic distances to dominant clades compared with VaxSeer (A/H3N2: 2.98 ± 2.13 *vs*. 5.89 ± 5.94, *P* = 0.1615; A/H1N1(pdm09): 1.01 ± 0.56 vs. 1.45 ± 0.38, *P* < 0.01) (Fig. 4C, D). It should be noted that vaccine production factors, such as viral stability in production systems, viral growth efficiency, and technical feasibility of recombinant have not been considered in our computational process. Nevertheless, our method helps to recommend vaccine strains antigenically similar to dominant variants.

**Fig. 4.**
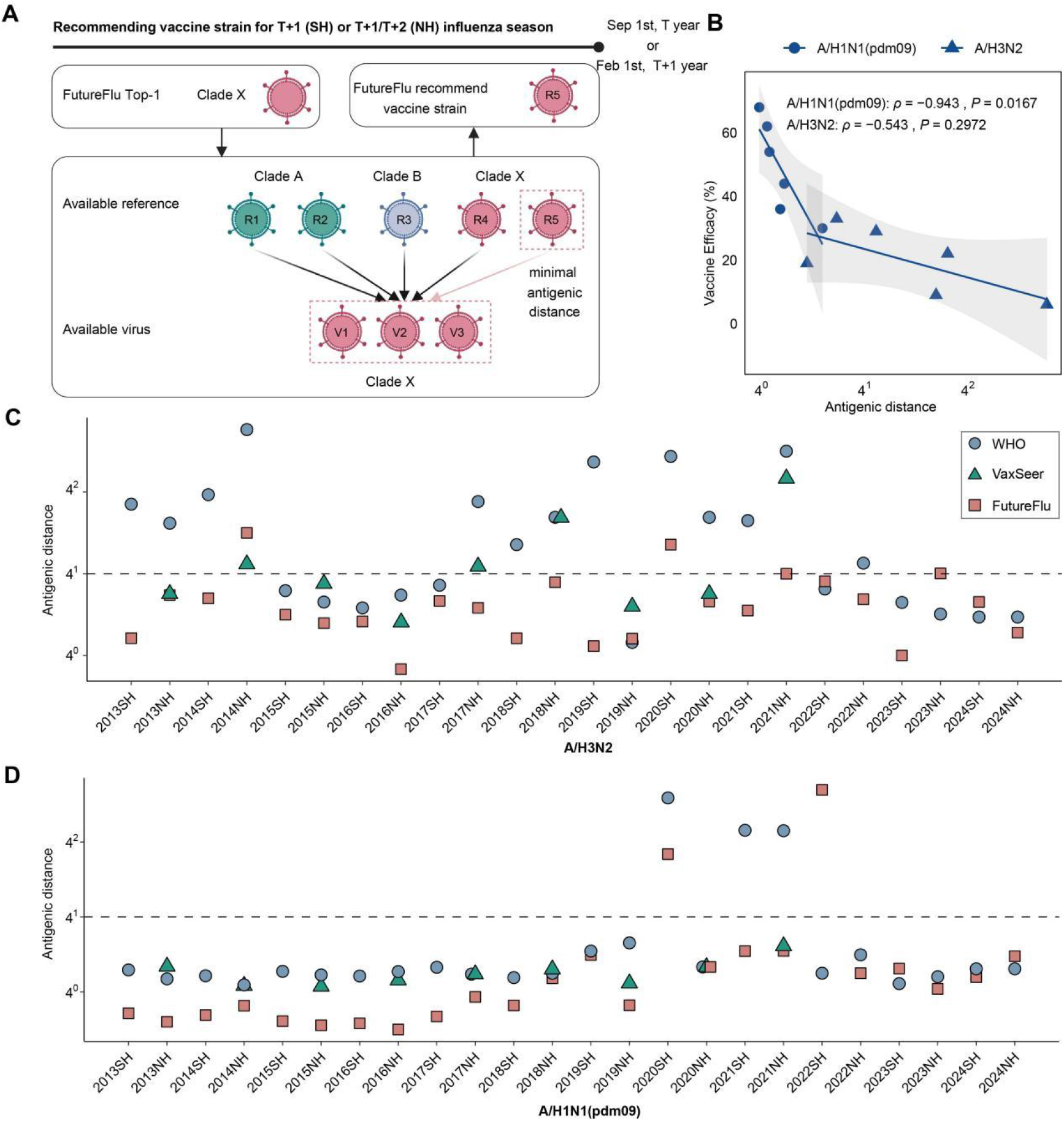
FutureFlu recommends vaccine strains antigenically similar to dominant variants. (**A**) FutureFlu’s vaccine strain recommendation process for upcoming influenza seasons. The cultured strain with minimal antigenic distance to the predicted Top-1 clade (clade X with the highest fitness score) is selected as the vaccine strain (R5). In HI assay, cultured strains (termed as “references”) are used to immunize ferrets for antisera production, while viral strains that react with these antisera are termed as “viruses”. Only references and viruses available before Sep 1^st^, T year (for SH) or Feb 1^st^, T+1 year (for NH) are considered in the selection process. (**B**) Correlation between antigenic distance and vaccine efficacy, with circles representing A/H1N1(pdm09) and triangles representing A/H3N2. (**C**) Antigenic distances between vaccine strains and dominant clade strains for A/H3N2 subtype. Vaccine strains are shown as blue circles for WHO (24 seasons), red squares for FutureFlu (24 seasons), and green triangles for VaxSeer (9 seasons from 2013NH to 2021NH, as reported in their publication). Antigenic distances below 4 indicate similarity, while antigenic distances ≥4 indicate dissimilarity. (**D**) Antigenic distances between vaccine strains and dominant clade strains for A/H1N1(pdm09) subtype. Panel (**A**) is created in BioRender.com.

Since 2024, A/H3N2 has been dominated by clade 3C.2a1b.2a.2a.3a.1, A/H1N1(pdm09) by 6B.1A.5a.2a, and B/Victoria by V1A.3a.2. These clades have evolved into multiple subclades, and we conducted our predictions at the subclade level. Based on sequence data submitted to GISAID before August 31^st^, 2025, FutureFlu predicted that the J.2 subclade carrying 24D, 25G, 112I, and 239I, D.3.1 subclade, and C.5.6.1 subclade showed the highest fitness to become dominant in the 2026SH season for A/H3N2, A/H1N1(pdm09), and B/Victoria, respectively, while several other subclades also had potential risk (fig. S10A). Detailed results can be viewed on our online recommendation platform of seasonal influenza vaccine strains (futureflu.com.cn). HI assay showed that J.2:24D,25G,112I,239I, J.2, J.2.3, and J.2.4 in A/H3N2 are antigenically distinct from each other, but the cultured strain A/Switzerland/47775/2024 in J.2.1 demonstrates broad cross-protection against these subclades. For the other subtypes, D.3.1, C.1.9.3, D.3, and C.1.9 of A/H1N1(pdm09) and C.5.6.1, C.5.6, C.5.7, C.5.1, and C.5 of B/Victoria are antigenically similar within their respective subtypes (fig. S10B). These results suggest that multiple antigenically distinct subclades of A/H3N2 will co-circulate in the 2026SH season, while multiple antigenically similar subclades of A/H1N1(pdm09) and B/Victoria will co-circulate.

## Discussion

The genetic diversity during influenza virus evolution leads to multiple variants co-circulating during the same period. Predicting the variant with best fitness that will be predominant in the upcoming season is both crucial and challenging for vaccine strain selection. Two kinds of methods are currently used to assess virus fitness. One strategy explicitly defines evolutionary hypotheses to predict properties of future viral populations based on past and present data. For instance, Łuksza et al. proposed a fitness model that uses genetic clades as basic units of prediction, establishing an approximate mapping between HA sequences and viral fitness(*11*). This model assigns positive effects to mutations in epitope sites to reflect cross-immunity properties, while potentially destabilizing mutations outside epitope regions are assigned negative effects, emphasizing the evolutionary trade-off between acquiring novel traits and maintaining basic functions. The second approach does not explicitly model viral fitness but rather predicts clade or strain frequencies based on genealogical tree data. For example, Neher et al. evaluate recent clade growth through local tree topology to identify high-fitness clades(*7*), while Hayati et al. combine phylogenetic tree reconstruction patterns with support vector machines to predict potentially circulating strains(*8*). More recently, Wang et al. developed a site-based dynamic model for mutation fitness prediction using linear extrapolation(*10*).

Beyond focusing on fitness modelling, several studies have developed models such as PREDAC(*32-34*) and VaxSeer(*31*) to predict the degree of cross-reactivity between any pair of strains, where hemagglutination inhibition (HI) assay data is essential. However, integrating HI data into prediction models presents challenges. Some A/H3N2 strains no longer react in HI assays, leading to increased use of neutralization assays to complement HI titer data(*35*). Additionally, the accuracy remains limited, which is why WHO has employed post-vaccination human serum panels to confirm the antigenic properties of circulating viruses for many years(*36*). Although these different types of serological data can be integrated using mixed analysis methods, it’s important to note that the interpretation of antigenic data involves considerable complexity.

Here, we developed FutureFlu, a computational framework that quantifies viral fitness by integrating three levels of evolutionary processes from variant emergence to dominance: genetic divergence at the molecular level, immune escape at the individual level, and transmission at the population level, using HA sequences and sero-positivity rate as input(*1, 3*). Higher fitness scores indicate greater potential for dominant circulation in upcoming influenza seasons. The retrospective analyses of three seasonal influenza viruses across 24 influenza seasons revealed significant correlation between clade fitness score and its observed frequency. FutureFlu supports two practical applications for vaccine strain recommendation: (1) prioritizing predicted dominant variants when antigenic assay data are unavailable, and (2) selecting cultured strains that are antigenically closest to predicted dominant variants when such assay data exist. Among the limited cultured strains, FutureFlu recommended antigenically similar vaccine strains for more influenza seasons, achieving smaller antigenic distances between vaccine and dominant strains compared to WHO and VaxSeer recommendations. While this study has not yet considered vaccine production factors(*37*), including adaptation stability(*38*), viral growth efficiency(*39*), and recombinant technical feasibility(*40*), our method effectively guides both vaccine strain recommendations and antigenic measurement strategies, such as conducting more intensive antigenic measurements for high-fitness clades.

To support ongoing seasonal influenza vaccine strain recommendations, we developed a vaccine strain recommendation platform (www.futureflu.com.cn). It will release prediction results before annual vaccine strain recommendation meetings, including three components, “Surveillance Dynamics of the Current Influenza Season”, “Potential Dominant Clade Ranking for the Next Influenza Season”, and “FutureFlu Recommends Vaccine Strains for the Next Influenza Season”. Using only sequence and epidemiological data, our model enables frequent updates of predictions, guiding virus isolation and early validation of risk mutations. While existing predictive models primarily focus on global-scale analysis, influenza virus evolution and circulation patterns exhibit distinct regional characteristics(*13, 41*). FutureFlu can provide region-specific vaccine recommendations based on local circulation patterns.

FutureFlu is a modular and probabilistic framework that integrates viral evolution across multiple scales. The modular design ensures both interpretability and extensibility for more components. Currently, advances in deep learning for mutation effects prediction(*18, 42*) will enhance fitness quantification by incorporating more accurate predictive models into each module. Given its biologically-grounded and modular design, this framework can be extended to address evolutionary prediction challenges in more respiratory viruses like SARS-CoV-2(*43*) and norovirus(*44*), guiding infectious disease prevention and control.

At present, a temporal limitation is evident: FutureFlu, like other existing models, can only assess the fitness potential of observed variants rather than predict the emergence of novel variants(*6*). In cases where predictions failed, about 50% of seasons were dominated by previously unobserved variants, highlighting the importance for emerging variant detection. Besides, DMS can systematically measure multiple mutation effects, such as replication efficiency(*12*), immune escape(*21*), and infectivity(*45*). Integrating these functional measurements into the prediction framework would enable identification of potentially circulating variants before they emerge in surveillance(*46*).

In summary, we proposed a theoretical framework that integrates molecular genetic divergence, individual immune escape, and population-scale transmission dynamics into a unified, quantitative measure of viral fitness. Applied to seasonal influenza virus as FutureFlu, this framework predicts dominant variants in upcoming seasons and supports evidence-based vaccine strain selection. Validation shows FutureFlu recommends antigenically matched vaccine strains more frequently than annual recommendations for A/H3N2, A/H1N1(pdm09), and B/Victoria subtypes. It is a proof for quantifying viral fitness landscape through integrating evolutionary dynamics from microscopic to macroscopic scales to improve vaccine strain selection and pandemic preparedness, enabling adaptation to other respiratory viruses.

## Acknowledgments

We gratefully acknowledge the authors from the originating and submitting laboratories where genetic sequence data were generated and shared via GISAID, enabling this research. We acknowledge High-performance Computing Platform and NCTIB Fund for R&D Platform for Cell and Gene Therapy of Suzhou Institute of Systems Medicine, Chinese Academy of Medical Sciences & Peking Union Medical College.

## Funding

Prevention and Control of Emerging and Major Infectious Diseases-National Science and Technology Major Project (2025ZD01901804)

CAMS Innovation Fund for Medical Sciences (CIFMS) (2021-I2M-1-061) CAMS Innovation Fund for Medical Sciences (CIFMS) (2022-I2M-1-021) CAMS Innovation Fund for Medical Sciences (CIFMS) (2022-I2M-3-001) China Postdoctoral Science Foundation (2025M772719)

Major Project of Guangzhou National Laboratory (GZNL2024A01015) National Natural Science Foundation of China (32370703)

National Natural Science Foundation of China (32500551)

Suzhou Applied Basic Research Program (General Program in Medical and Health Sciences) (SYW2024065)

General Program of National Natural Science Foundation of China (82372225)

Science and Technology Leading Talent Program of Yunnan Province (202405AB350002) Yunnan Provincial Major Science and Technology Projects (202402AA310024).

## Author contributions

Conceptualization: A.W., Y.W., D.W., T.J., J.W.

Methodology: L.Q., X.L.

Investigation: L.Q., L.Y., Y.Q., J.L., W.X.

Validation: L.Q., M.Z., X.L., Y.M., Z.L., Z.L.

Visualization: L.Q., A.W., X.L., J.Y., H.W.

Funding acquisition: A.W., Y.W., X.D., H.Z., L.Q., Q.L.

Project administration: A.W., Y.W.

Supervision: A.W., Y.W., D.W., T.J., J.W

Writing – original draft: L.Q., X.D.

Writing – review & editing: A.W., Y.W., D.W., T.J., J.W., L.Q., X.D., M.Z., Y.Q., X.L.

## Data and materials availability

All influenza HA sequences and their metadata are available through https://gisaid.org/. The accession IDs for the HA proteins used in this study are available in our GitHub repository at https://github.com/wuaipinglab/FutureFlu. The HI results are collected from WHO’s annual influenza vaccine composition consultation documents published between 2003 and September 2025(https://www.crick.ac.uk/research/platforms-and-facilities/worldwide-influenza-centre/annual-and-interim-reports). The human influenza vaccine composition data are obtained from https://gisaid.org/resources/human-influenza-vaccine-composition/. We used the following Protein Data Bank (PDB) identifiers: 4fnk, 3lzg, and 4fqm. The model code is available at https://github.com/wuaipinglab/FutureFlu.

## References and Notes

1. V. N. Petrova, C. A. Russell, The evolution of seasonal influenza viruses. Nat Rev Microbiol 16, 47–60 (2018).

2. M. I. Nelson, E. C. Holmes, The evolution of epidemic influenza. Nat Rev Genet 8, 196–205 (2007).

3. I. M. Rouzine, G. Rozhnova, Antigenic evolution of viruses in host populations. PLoS Pathog 14, e1007291 (2018).

4. A. M. Carabelli et al., SARS-CoV-2 variant biology: immune escape, transmission and fitness. Nat Rev Microbiol 21, 162–177 (2023).

5. M. Meijers et al., Concepts and Methods for Predicting Viral Evolution. Methods Mol Biol 2890, 253–290 (2025).

6. D. H. Morris et al., Predictive Modeling of Influenza Shows the Promise of Applied Evolutionary Biology. Trends Microbiol 26, 102–118 (2018).

7. R. A. Neher, C. A. Russell, B. I. Shraiman, Predicting evolution from the shape of genealogical trees. Elife 3, (2014).

8. M. Hayati, B. Sobkowiak, J. E. Stockdale, C. Colijn, Phylogenetic identification of influenza virus candidates for seasonal vaccines. Sci Adv 9, eabp9185 (2023).

9. J. Huddleston et al., Integrating genotypes and phenotypes improves long-term forecasts of seasonal influenza A/H3N2 evolution. Elife 9, (2020).

10. J. Lou et al., Predictive evolutionary modelling for influenza virus by site-based dynamics of mutations. Nat Commun 15, 2546 (2024).

11. M. Luksza, M. Lässig, A predictive fitness model for influenza. Nature 507, 57–61 (2014).

12. J. M. Lee et al., Deep mutational scanning of hemagglutinin helps predict evolutionary fates of human H3N2 influenza variants. Proc Natl Acad Sci U S A 115, E8276–e8285 (2018).

13. T. Bedford et al., Global circulation patterns of seasonal influenza viruses vary with antigenic drift. Nature 523, 217–220 (2015).

14. A. C. Shih, T. C. Hsiao, M. S. Ho, W. H. Li, Simultaneous amino acid substitutions at antigenic sites drive influenza A hemagglutinin evolution. Proc Natl Acad Sci U S A 104, 6283–6288 (2007).

15. B. F. Koel et al., Substitutions near the receptor binding site determine major antigenic change during influenza virus evolution. Science 342, 976–979 (2013).

16. M. Zhang et al., Retrospective immunogenicity analysis of seasonal flu H3N2 vaccines recommended in the past ten years using immunized animal sera. EBioMedicine 86, 104350 (2022).

17. W. Han et al., Predicting the antigenic evolution of SARS-COV-2 with deep learning. Nat Commun 14, 3478 (2023).

18. Z. Nie et al., A unified evolution-driven deep learning framework for virus variation driver prediction. Nature Machine Intelligence 7, 131–144 (2025).

19. A. J. Riesselman, J. B. Ingraham, D. S. Marks, Deep generative models of genetic variation capture the effects of mutations. Nat Methods 15, 816–822 (2018).

20. N. N. Thadani et al., Learning from prepandemic data to forecast viral escape. Nature 622, 818–825 (2023).

21. J. M. Taft et al., Deep mutational learning predicts ACE2 binding and antibody escape to combinatorial mutations in the SARS-CoV-2 receptor-binding domain. Cell 185, 4008–4022.e4014 (2022).

22. B. S. Ho, K. M. Chao, Data-driven interdisciplinary mathematical modelling quantitatively unveils competition dynamics of co-circulating influenza strains. J Transl Med 15, 163 (2017).

23. S. P. J. de Jong, A. J. K. Conlan, A. X. Han, C. A. Russell, Competition between transmission lineages mediated by human mobility shapes seasonal influenza epidemics in the US. Nat Commun 16, 4605 (2025).

24. T. A. Hopf et al., Mutation effects predicted from sequence co-variation. Nat Biotechnol 35, 128–135 (2017).

25. L. C. Martin, G. B. Gloor, S. D. Dunn, L. M. Wahl, Using information theory to search for co-evolving residues in proteins. Bioinformatics 21, 4116–4124 (2005).

26. D. de Juan, F. Pazos, A. Valencia, Emerging methods in protein co-evolution. Nature Reviews Genetics 14, 249–261 (2013).

27. J. Lou et al., Predicting the dominant influenza A serotype by quantifying mutation activities. Int J Infect Dis 100, 255–257 (2020).

28. S. Elbe, G. Buckland-Merrett, Data, disease and diplomacy: GISAID’s innovative contribution to global health. Glob Chall 1, 33–46 (2017).

29. I. Khandaker et al., Molecular evolution of the hemagglutinin and neuraminidase genes of pandemic (H1N1) 2009 influenza viruses in Sendai, Japan, during 2009-2011. Virus Genes 47, 456–466 (2013).

30. M. Cardenas et al., Amino acid 138 in the HA of a H3N2 subtype influenza A virus increases affinity for the lower respiratory tract and alveolar macrophages in pigs. PLoS Pathog 20, e1012026 (2024).

31. W. Shi, J. Wohlwend, M. Wu, R. Barzilay, Influenza vaccine strain selection with an AI-based evolutionary and antigenicity model. Nat Med, (2025).

32. X. Du et al., Mapping of H3N2 influenza antigenic evolution in China reveals a strategy for vaccine strain recommendation. Nat Commun 3, 709 (2012).

33. J. Meng et al., PREDAC-CNN: predicting antigenic clusters of seasonal influenza A viruses with convolutional neural network. Brief Bioinform 25, (2024).

34. Y. Peng et al., Automated recommendation of the seasonal influenza vaccine strain with PREDAC. Biosafety and Health 2, 117–119 (2020).

35. Y. Lin et al., The characteristics and antigenic properties of recently emerged subclade 3C.3a and 3C.2a human influenza A(H3N2) viruses passaged in MDCK cells. Influenza Other Respir Viruses 11, 263–274 (2017).

36. J. M. Fonville et al., Antigenic Maps of Influenza A(H3N2) Produced With Human Antisera Obtained After Primary Infection. J Infect Dis 213, 31–38 (2016).

37. P. C. Soema, R. Kompier, J. P. Amorij, G. F. Kersten, Current and next generation influenza vaccines: Formulation and production strategies. Eur J Pharm Biopharm 94, 251–263 (2015).

38. D. M. Skowronski et al., Low 2012-13 influenza vaccine effectiveness associated with mutation in the egg-adapted H3N2 vaccine strain not antigenic drift in circulating viruses. PLoS One 9, e92153 (2014).

39. M. Nemoto et al., Growth properties of recombinant equine influenza viruses with different backbones generated by reverse genetics in embryonated chicken eggs. Arch Virol 170, 181 (2025).

40. R. E. Colombo et al., Randomized pragmatic trial of the comparative effectiveness of chicken egg-based inactivated, mammalian cell-culture-based inactivated, and recombinant protein quadrivalent seasonal influenza vaccines in United States Military Health System beneficiaries. Clin Infect Dis, (2025).

41. J. Tamerius et al., Global influenza seasonality: reconciling patterns across temperate and tropical regions. Environ Health Perspect 119, 439–445 (2011).

42. Y.-F. Pan et al., Predicting the Evolutionary and Functional Landscapes of Viruses with a Unified Nucleotide-Protein Language Model: LucaVirus. bioRxiv, 2025.2006.2014.659722 (2025).

43. H. Ren et al., Early warning of emerging infectious diseases based on multimodal data. Biosafety and Health 5, 193–203 (2023).

44. B. V. V. Prasad et al., Norovirus replication, host interactions and vaccine advances. Nature Reviews Microbiology 23, 385–401 (2025).

45. B. Dadonaite et al., Deep mutational scanning of H5 hemagglutinin to inform influenza virus surveillance. PLoS Biol 22, e3002916 (2024).

46. M. B. Doud, J. D. Bloom, Accurate Measurement of the Effects of All Amino-Acid Mutations on Influenza Hemagglutinin. Viruses 8, (2016).

47. K. Katoh, D. M. Standley, MAFFT multiple sequence alignment software version 7: improvements in performance and usability. Mol Biol Evol 30, 772–780 (2013).

48. M. H. Wang et al., Characterization of key amino acid substitutions and dynamics of the influenza virus H3N2 hemagglutinin. J Infect 83, 671–677 (2021).

49. I. Aksamentov, C. Roemer, E. Hodcroft, R. Neher, Nextclade: clade assignment, mutation calling and quality control for viral genomes. Journal of Open Source Software 6, 3773 (2021).

50. J. Hadfield et al., Nextstrain: real-time tracking of pathogen evolution. Bioinformatics 34, 4121–4123 (2018).

51. J. C. Pedersen, Hemagglutination-inhibition assay for influenza virus subtype identification and the detection and quantitation of serum antibodies to influenza virus. Methods Mol Biol 1161, 11–25 (2014).

